# Genetic evolution analysis of hemagglutinin and neuraminidase genes of H9N2 avian influenza virus in external environment of some areas of Yunnan Province, China from 2020 to 2023

**DOI:** 10.1101/2024.02.24.581849

**Authors:** Zhaosheng Liu, Xiaoqing Fu, Yaoyao Chen, Yanhong Sun, Meiling Zhang, Xiaoyu Han, Xiaonan Zhao, Jienan Zhou

## Abstract

**Objective:** To understand the molecular characteristics and genetic variation of hemagglutinin (HA) and neuraminidase (NA) genes of H9N2 subtype avian influenza virus (AIV) in the external environment of Yunnan Province, and to provide evidence for the prevention and control of H9N2 subtype AIV in this area.

**Results:** The HA and NA genes of 20 isolates belonged to Y280-like sub-branch. The nucleotide and amino acid homology of HA gene were 88.46%-99.81% and 89.08% -99.61%, respectively. The nucleotide and amino acid homology of NA gene were 88.85%-100% and 90.09%-100%, respectively. The HA protein cleavage site of 20 isolates was PSRSSRGLF, which was consistent with the molecular characteristics of low pathogenic avian influenza virus. The 226 th and 228 th positions of the receptor binding site are all L and G, which have the ability to bind to the mammalian sialic acid α-2,6 sialic acid receptor; HA protein had 7-8 glycosylation sites, and the main variation was the deletion of one site at 218 and the addition of one glycosylation site at 313 and 491. The main antigenic sites were G90E, S/T145D, D/N153G, A/S/F168N/E, T200R, N/D201G/T mutations. The NA protein neck of 20 isolates lacked 3 amino acids (TEI), which had the molecular characteristics of highly pathogenic avian influenza. NA protein had 6-8 glycosylation sites. The main variation was that two isolates increased a new glycosylation site NPTQ at the 2nd position, and one isolate increased a new glycosylation site NTTI at the 67th position. All isolates lost one site at the 402nd position, and some isolates lost at the 83rd and 365th positions. The amino acids at the active site and key site of NA protease were not mutated, and the isolates did not produce resistance to neuraminidase inhibitors.

**Conclusion:** The HA and NA genes of H9N2 subtype avian influenza virus in Yunnan Province have evolved continuously, but they still belong to the Y280 branch of the Eurasian lineage. Mutations in key sites may cause increased infectivity and transmission. The monitoring of H9N2 subtype avian influenza virus should be strengthened to prevent it from breaking the interspecies barrier and spreading to humans and lower mammals, so as to prevent the outbreak of H9N2 subtype avian influenza.

---

Avian Influenza (AI) is an avian viral infectious disease caused by influenza A virus, which is listed as a notifiable animal infectious disease by the International Veterinary Agency (OIE). H9N2 subtype avian influenza virus (AIV) is widely distributed all over the world. It was first isolated in North America in 1966 and first discovered in China in 1994[1]. For the first time, the continuous spread of H9N2 subtype AIV was found, which brought huge economic losses to China ‘s poultry industry. Since 1998, H9N2 subtype AIVs have been isolated from pigs and humans[2].In 1997,1999 and 2004-2005, there were outbreaks of H5N1 and H9N2 subtype avian influenza in Hong Kong, Thailand and Vietnam, and even cases of virus breaking through the interspecies barrier to infect humans. In 2013, human infection with H7N9 and H10N8 subtype avian influenza[3, 4] appeared in China, giving avian influenza a new public health significance. It not only puts pressure on the poultry industry economy, but also poses a potential threat to human public health.

The genome of avian influenza virus is composed of single-strand negative-strand RNA, which is divided into 8 gene fragments and encodes 10 genes. HA and NA are the main surface antigens, and there are 16 HA subtypes and 9 NA subtypes[5]. The cleavage site and glycosylation site of HA are the key factors affecting pathogenicity and virulence, and glycosylation affects the binding level of virus to host cells. Avian influenza is divided into non-pathogenic avian influenza (NPAI), low pathogenic avian influenza (LPAI) and highly pathogenic avian influenza (HPAI). In China, H9N2 subtype mainly causes LPAI.Although the mortality rate is not high, it has a significant impact on the growth, immune and reproductive systems of poultry, which can easily lead to mixed infection and cause more serious economic losses[6]. At present, vaccination is the main means of prevention and control of avian influenza, but the continuous variation of the virus increases the difficulty of prevention and control, especially the H9N2 subtype AIV, whose genetic evolution characteristics pose a potential threat to the poultry industry and human health. Real-time monitoring of the prevalence of avian influenza and understanding of its genetic evolution characteristics are crucial for the scientific development of prevention and control strategies.

In summary, avian influenza is not only a challenge for the poultry industry, but also a complex problem involving public health, economy and environment. Strengthening the monitoring and research of H9N2 subtype AIV and formulating scientific prevention and control strategies will help maintain the sustainable development of the poultry industry and effectively protect public health.

## 1. Materials and Methods

### 1.1 Materials

#### 1.1.1 Virus and chicken embryo

The 20 strains of H9N2 subtype AIV used in this experiment were isolated from environmental specimens collected from live poultry markets in the external environment avian influenza monitoring mission of Yunnan Province from 2020 to 2023 (cage surface swab specimens, sewage for cleaning poultry, feces specimens, poultry drinking water, swab specimens on the surface of slaughtered or placed poultry meat boards). The specific information is shown in Table 1. After H9N2 subtype was detected by real-time fluorescence quantitative PCR (Real time-PCR), it was obtained by inoculating chicken embryo culture. SPF chicken embryos, 9∼10 days of age, purchased from Shandong Fapasi company.

**Table 1.** The basic situation of 20 H9N2 subtype AIV isolates.

| Number | Strain | Abbreviation | City | Year |
| --- | --- | --- | --- | --- |
| 1 | A/Enviroment/Yunnan/KM061/2021 | KM061 | kunming yunnan China | 2021 |
| 2 | A/Enviroment/Yunnan/KM029/2020 | KM029 | kunming yunnan China | 2020 |
| 3 | A/Enviroment/Yunnan/KM034/2022 | KM034 | Kunming yunnan China | 2022 |
| 4 | A/Enviroment/Yunnan/DQ057/2022 | DQ057 | Diqing Yunnan China | 2022 |
| 5 | A/Enviroment/Yunnan/DQ073/2022 | DQ073 | Diqing Yunnan China | 2022 |
| 6 | A/Enviroment/Yunnan/CX007/2022 | CX007 | Chuxiong Yunnan China | 2022 |
| 7 | A/Enviroment/Yunnan/CX022/2022 | CX022 | Chuxiong Yunnan China | 2022 |
| 8 | A/Enviroment/Yunnan/HH085/2021 | HH085 | Red River Yunnan China | 2021 |
| 9 | A/Enviroment/Yunnan/CX034/2021 | CX034 | Chuxiong Yunnan China | 2021 |
| 10 | A/Enviroment/Yunnan/KM038/2023 | KM038 | Kunming yunnan China | 2023 |
| 11 | A/Enviroment/Yunnan/PE036/2023 | PE036 | Pu 'er Yunnan China | 2023 |
| 12 | A/Enviroment/Yunnan/DH084/2021 | DH084 | Dehong Yunnan China | 2021 |
| 13 | A/Enviroment/Yunnan/DH038/2020 | DH038 | Dehong Yunnan China | 2020 |
| 14 | A/Enviroment/Yunnan/NJ019/2021 | NJ019 | Nujiang Yunnan China | 2021 |
| 15 | A/Enviroment/Yunnan/NJ014/2020 | NJ014 | Nujiang Yunnan China | 2020 |
| 16 | A/Enviroment/Yunnan/BS013/2023 | BS013 | Baoshan Yunnan China | 2023 |
| 17 | A/Enviroment/Yunnan/YX023/2021 | YX023 | Yuxi Yunnan China | 2021 |
| 18 | A/Enviroment/Yunnan/CX023/2023 | CX023 | Chuxiong, Yunnan China | 2023 |
| 19 | A/Enviroment/Yunnan/YX081/2020 | YX081 | Yuxi Yunnan China | 2020 |
| 20 | A/Enviroment/Yunnan/CX021/2020 | CX021 | Chuxiong, Yunnan China | 2020 |

#### 1.1.2 Main reagents

Nucleic acid extraction reagent was purchased from Xi ‘an Tianlong Technology Co., Ltd. (magnetic bead method, item number : 221201DOT183) ; avian influenza virus H9N2 subtype nucleic acid detection kit (fluorescence quantitative PCR) was purchased from Beijing Zhuocheng Huisheng Biotechnology Co., Ltd. ; one Step RT-PCR Kit purchased from ABI ; the capture amplification primers were referred to the ‘National Influenza Monitoring Technology Guide (2017 Edition)’, and the specific primers were synthesized by Beijing Qingke Biotechnology Co., Ltd. ; nextera XT DNA Sample Preparation Kit, Nextera XT Index Kit and Miseq Sequencing Kit (MiSeq Reagent Kits v2,300 cylce) were purchased from Illumina.

### 1.2 Methods

#### 1.2.1 Virus isolation

The positive swab samples were detected by H9 subtype influenza virus fluorescent RT-PCR detection kit and influenza virus N2 subtype fluorescent RT-PCR detection kit, centrifuged at 8000 r / min for 5 min, and 0.40 mL of the supernatant was inoculated into 10-day-old chicken embryos. After 72h of incubation in a 37 °C incubator, the allantoic fluid of chicken embryos was collected, and the chicken embryos that died within 24h were discarded for serum identification test. The H9 subtype was verified again by fluorescence RT-PCR, and the H9 positive strain was determined to be stored at-80 °C.

#### 1.2.2 Nucleic acid extraction and amplification

Nucleic acid extraction was carried out according to the instructions of Tianlong nucleic acid purification reagent. Nucleic acid samples were frozen in a refrigerator at -80 °C for later use. The full-length HA and NA genes were amplified according to the instructions of One Step RT-PCR Kit.

#### 1.2.3 HA, NA gene sequence analysis

The sequencing data were processed by CLC genomics analysis software, and the homology analysis was carried out by Lasergene Meg Align software. The Neighbor-Joining method in MEGA7.0 software was used to construct the genetic evolution tree of HA and NA genes. The calculation method was Maximum Composite Likelihood, and the bootsrap value was 1000. NetNGlyc 1.0 Server software was used to analyze the N-glycosylation sites of the amino acid sequence. The HA and NA gene sequences of H9N2 subtype avian influenza classical strains and vaccine strains were obtained from GenBank as the reference sequences for analysis and comparison: classical strains: A/chichen/Beijing/1994, A/chichen/Guangxi-55/2005, A/chichen/HongKong/G9/1997,A/chichen/Korea/MS96/96, A/duck/Hongkong/Y439/1997, A/duck/Hongkong/Y280/1997, A/Quail/Hongkong/G1/1997, A/turkey/Wisconsin/1/1966 vaccine strains:A/chicken/Jiangsu/WJ57/2012,A/chicken/Shandong/6/1996, A/chicken/Shanghai/F98/1998.

## 2. Results and analysis

### 2.1 HA gene of H9N2 subtype avian influenza virus

#### 2.1.1 Genetic evolution and homology analysis of HA gene

MEGA7.0 was used to analyze the genetic evolution of the HA gene of the reference sequence and the 20 H9N2 virus isolates obtained in this experiment, and the genetic evolution tree was drawn (Fig.1). The evolutionary tree of H9N2 subtype avian influenza virus was divided into two strains of Eurasia and North America. The Eurasian strain can be divided into three groups, namely Y280-like, Y439-like and G1-like branches. The analysis results show that the genetic distance between the HA genes of the 20 isolates is relatively close, and all belong to the Y280 branch of the Eurasian lineage.

**Figure 1.**
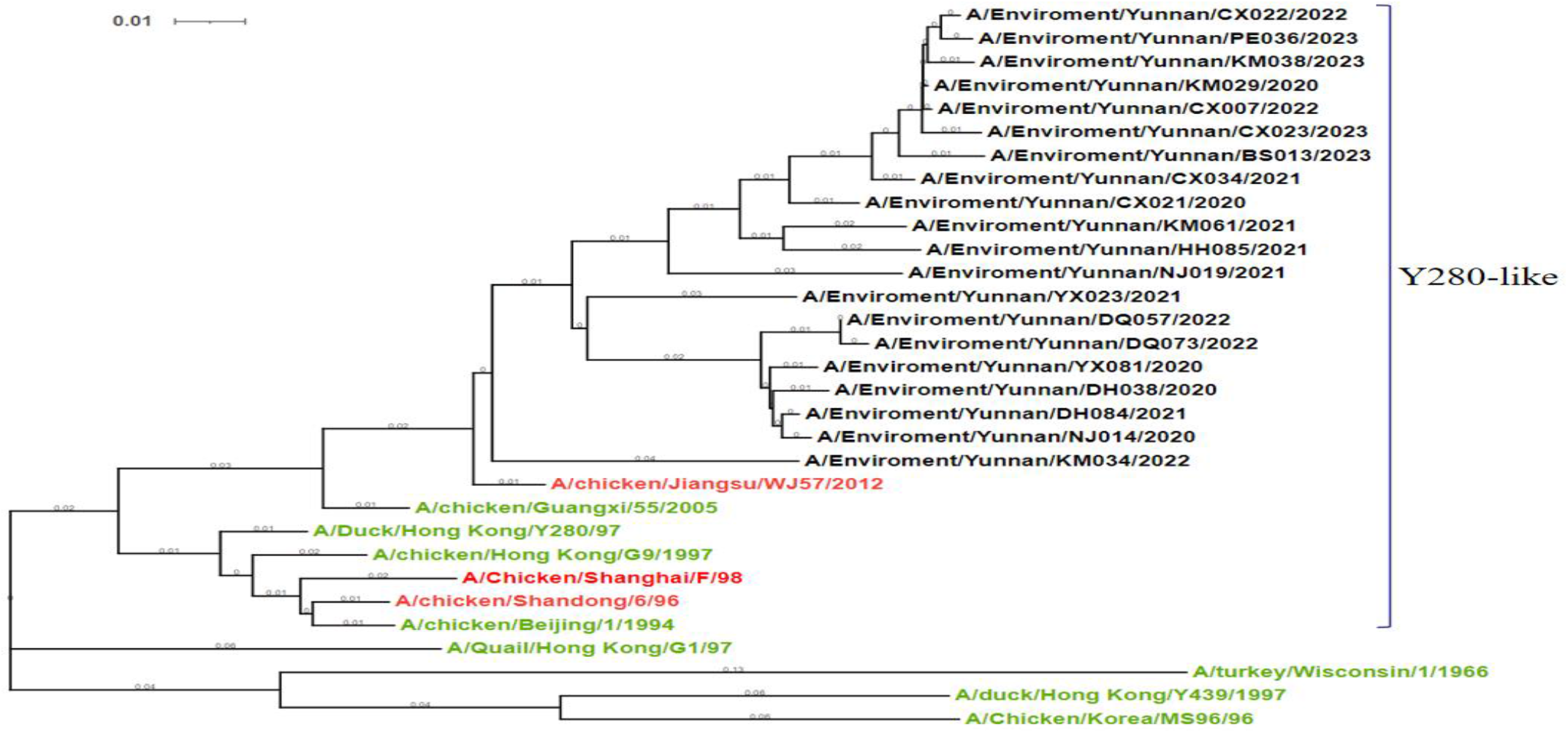
Phylogenetic tree of HA gene of 20 H9N2 subtype AIV isolates. **Note: Red represents vaccine strain; green represents the classic strain; black represents the isolate;**

**Figure 2.**
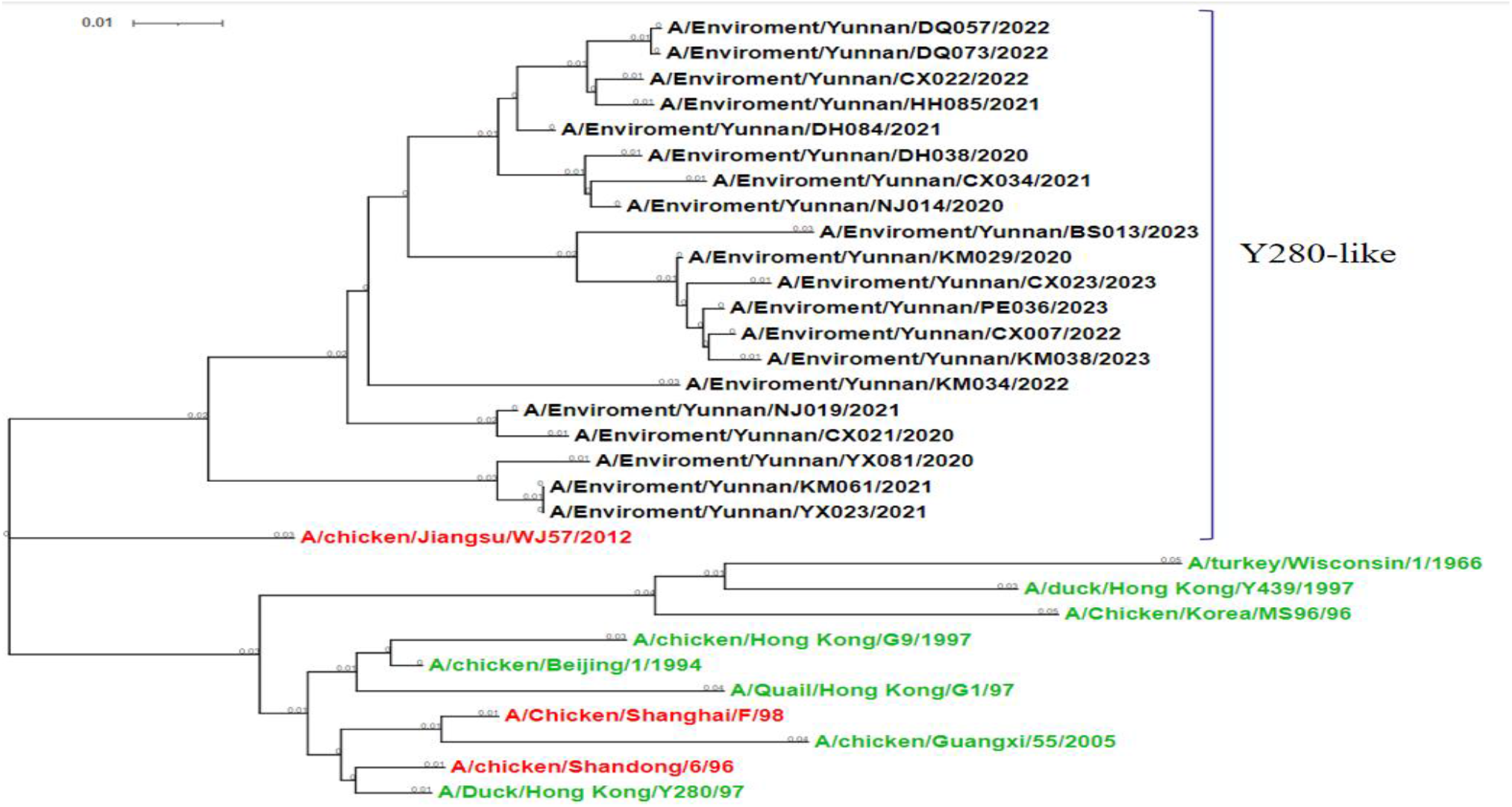
Phylogenetic tree of HA gene of 20 H9N2 subtype AIV isolates. **Note: Red represents vaccine strain; green represents the classic strain; black represents the isolate;**.

The nucleotide homology of HA gene among the 20 isolates was 88.46 % -99.81 %, and the deduced amino acid homology was 89.08 % -99.61 % (Table 2). Among them, the homology within the strain group in 2020 was 91.27 % -98.62 % and 91.63-98.64 %, respectively. The homology within the strain group in 2021 was 90.99 % -96.31 % and 91.35 % -96.37 %, respectively.

**Table 2.**
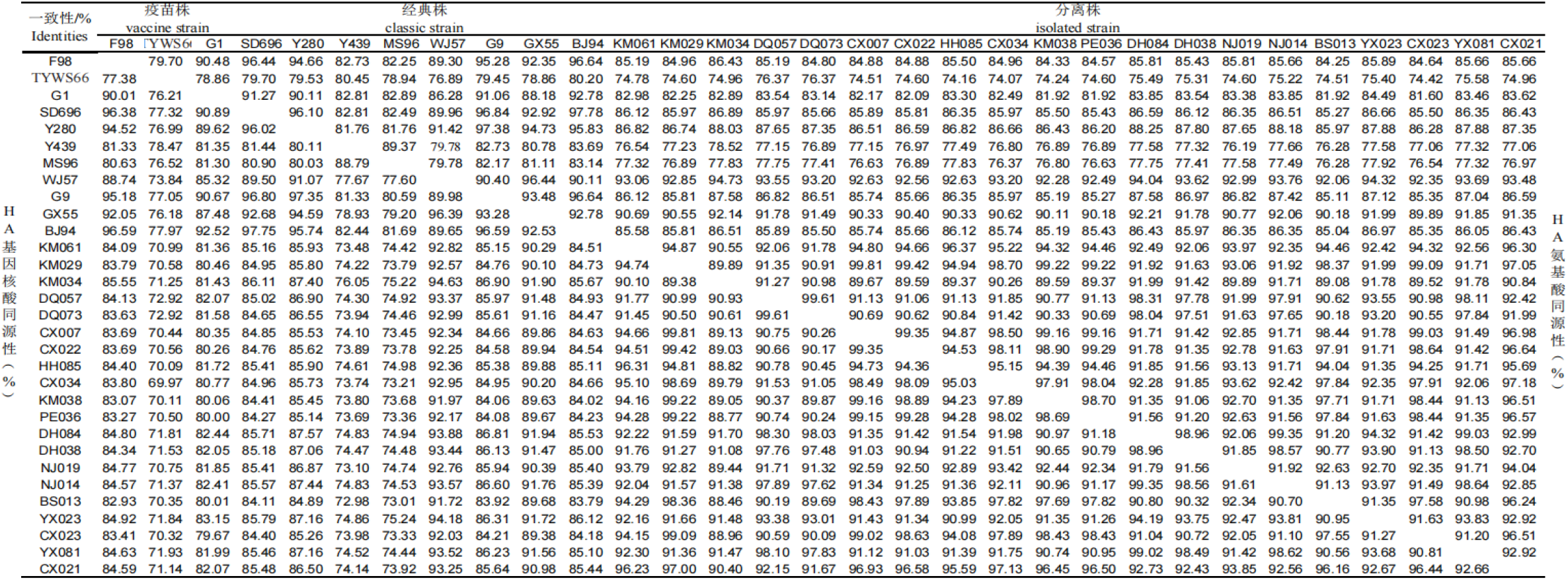
Comparison of nucleotide and amino acid sequences of HA genes of 20 H9N2 subtype AIV isolates.

The homology within the strain group in 2022 was 89.03 % -99.61 % and 89.59 % -99.61 %, respectively. The homology within the strain group in 2023 was 97.55 % -98.69 % and 97.58 % -98.70 %, respectively. In addition, the nucleotide and amino acid homology of the 20 strains with the vaccine strain WJ57 was the highest, which was 91.72 % -94.63 and 92.06 % -94.73 %, respectively. The homology with the vaccine strain F98 was the lowest, which was 82.93 % - 85.55 % and 84.25 % -86.43 %, respectively. The nucleotide and amino acid homology of the 20 strains was the highest with the classical strain GX55, which was 89.38 % -91.94 % and 89.89 % -92.21 %, respectively. The homology with the classical strain TYWS66 was the lowest, which was 69.97 % -72.92 % and 74.07 % -76.37 %, respectively.

#### 2.1.2 Analysis of key sites of HA gene

The complete ORF of HA gene sequence of 20 H9N2 subtype AIVs was composed of 1683 nucleotides, encoding 560 amino acids. The cleavage site of HA protein of 20 isolates was located at 333-341 amino acid sequence, and the sequence was PSRSSRGLF, which was consistent with the characteristics of low pathogenic avian influenza virus. Comparing the reference strains of the isolates, it was found that there were amino acid mutations at multiple sites in the HA protein, in which the amino acid at the 334th cleavage site was mutated from non-polar amino acid A to polar amino acid S;

The sialic acid receptor binding site at position 191 of HA was aspartic acid (N), which was consistent with the classical characteristics of avian influenza virus isolated in China. From 2020 to 2023, the isolates were relatively conserved at positions 109, 161, 202, 203, and 236, followed by Y, W, L, Y, and G, and there were variations at positions 163, 191, 198, and 199. Compared with the reference strain F98, a total of 11 isolates had a mutation T → N at position 163 from 2020 to 2023, including 2 isolates in 2020, 3 isolates in 2021, 2 isolates in 2022, and 4 isolates in 2023.

The 19 isolates had a mutation A → T (V) at position 198, including 1 strain 198V and 4 strains 198T in 2020, 2 strains 198V and 3 strains 198T in 2021, 2 strains 198T and 2 strains 198V in 2022, and 4 strains 198V in 2023. The 10 isolates had a mutation S → N at position 199, of which 3 were from 2020 to 2022 and 1 was from 2023. There was a mutation Q → L at position 234 in 20 isolates. There were mutations Q → L at position 234 and Q → M at position 235 in the left arm of the receptor binding site of 20 isolates. This type of mutation has the potential characteristics of binding to mammalian sialic acid α-2,6 receptor. There was a mutation K (R) → N (T) at the 149th site of the right arm of the 19 isolates, including 2 strains of 141 N and 3 strains of 149 T in 2020, 4 strains of 149 N and 2 strains of 149 T in 2021, 2 strains of 149 N and 2 strains of 149 T in 2022, and 4 strains of 149 N in 2023. There were 12 strains with mutation A → T (S) at the 150 th site of the right arm, including 2 strains of 150 T in 2020, 4 strains of 150 T in 2021, 2 strains of 150 T in 2022, 3 strains of 150 T and 1 strain of 150 S in 2023. The 7 antigenic sites of 20 isolates were analyzed. The results showed that the 80 th antigenic site of 19 strains in 2020-2023 was glutamic acid (E), which was consistent with the classical strain G1. Except that the 145th glycosylation site of KM034 isolate in 2023 was aspartic acid (D), the other 19 isolates had S/T145D mutation. The 20 isolates had different mutations at the 153 and 167 antigen sites. Although they showed diversity, they were consistent with the classical strains or vaccine strains. There were new amino acid mutations at the antigen sites of 168,200 and 201 in isolates from different years, which were A / S / F168E / N, T200R and D / N201T / G / Q (Table 3).

**Table 3.**
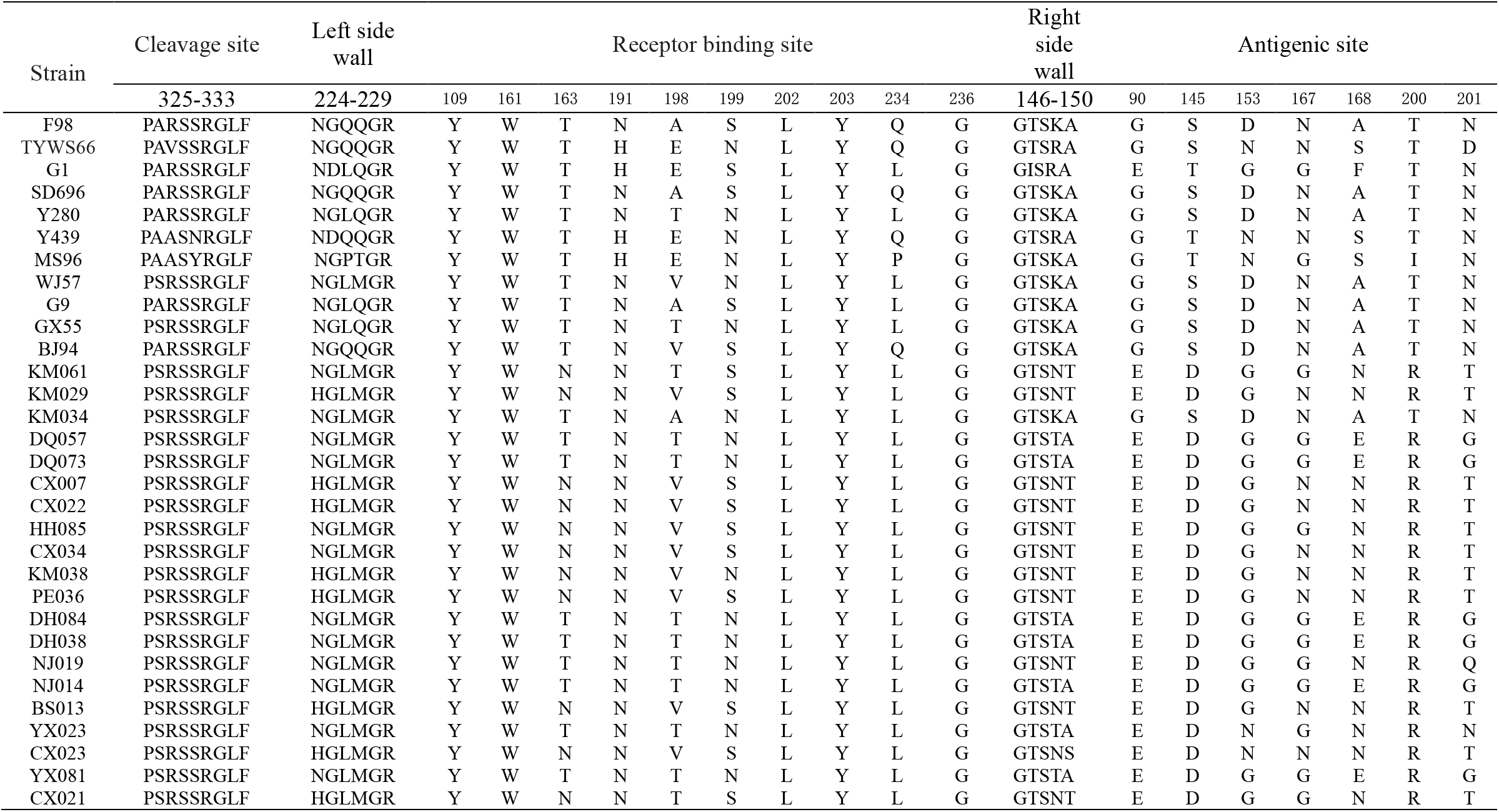
Analysis of HA protein receptor binding sites and antigenic sites of 20 H9N2 subtype AIV isolates.

The potential glycosylation sites of HA protein of 20 isolates were analyzed. The results showed that 20 isolates had 7-8 potential glycosylation sites. Among them, one isolate (NJ019) in 2021 had a change in the glycosylation site of NSTD at position 29, and the isolates in other years were relatively conservative at positions 29,82,141,298,305 and 313. Compared with the classical strain and the vaccine strain, the isolates had 10 glycosylation sites missing, and some isolates increased 2 glycosylation sites (491 and 550) (Table 4).

**Table 4.** Analysis of potential glycosylation sites of HA in 20 H9N2 subtype AIV isolates.

| Strain | Potential glycosylation sites of HA |  |  |  |  |  |  |  |  |  |  |  |
| --- | --- | --- | --- | --- | --- | --- | --- | --- | --- | --- | --- | --- |
|  | 29 | 82 | 137 | 141 | 218 | 298 | 305 | 313 | 491 | 492 | 550 | 551 |
| F98 | NSTE | NPSC | — | NVSY | NRTF | NTTL | NVSK | — | — | NGTY | — | NGSC |
| TYWS66 | NSTE | NPSC | — | NVTY | NRTF | NTTL | NISK | — | — | NGTY | — | NGSC |
| G1 | NSTE | NPSC | — | NVTY | NRTF | NSTL | NISK | — | — | NGTY | — | — |
| SD696 | — | — | NVSY | — | — | — | — | — | — | — | — | — |
| Y280 | NSTE | NPSC | — | NVSY | NRTF | NTTL | NVSK | — | — | NGTY | — | — |
| Y439 | NSTE | NPSC | — | NVTY | NRTF | NTTL | NVSK | — | — | NGTY | — | NGSC |
| MS96 | NSTE | NPSC | — | NVTY | NRTF | NTTL | NVSK | — | — | NGTY | — | NGSC |
| WJ57 | NSTE | NPSC | — | NVSY | — | NTTL | NVSK | NCSK | — | NGTY | — | NGSC |
| G9 | NSTE | NPSC | — | NVSY | NRTF | NTTL | NVSK | — | — | NGTY | — | NGSC |
| GX55 | NSTE | NPSC | — | NVSY | NRTF | NTTL | NVSK | NCSK | — | NGTY | — | NGSC |
| BJ94 | NSTE | NPSC | — | NVTY | NRTF | NTTL | NVSK | — | — | NGTY | — | NGSC |
| KM061 | NSTE | NPSC | — | NVSY | — | NTTL | NVSK | NCSK | — | NGTY | — | NGSC |
| KM029 | NSTE | NPSC | — | NVSY | — | NTTL | NVSK | NCSK | NGTY | — | NGSC | — |
| KM034 | NSTE | NPSC | — | NVSY | — | NTTL | NVSK | NCSK | — | NGTY | — | NGSC |
| DQ057 | NSTE | NPSC | — | NVSY | — | NTTL | NVSK | NCSK | — | NGTY | — | NGSC |
| DQ073 | NSTE | NPSC | — | NVSY | — | NTTL | NVSK | NCSK | — | NGTY | — | NGSC |
| CX007 | NSTE | NPSC | — | NVSY | — | NTTL | NVSK | NCSK | NGTY | — | NGSC | — |
| CX022 | NSTE | NPSC | — | NVSY | — | NTTL | NVSK | NCSK | NGTY | — | NGSC | — |
| HH085 | NSTE | NPSC | — | NVSY | — | NTTL | NVSK | NCSK | NGTY | — | NGSC | — |
| CX034 | NSTE | NPSC | — | NVSY | — | NTTL | NVSK | NCSK | NGTY | — | NGSC | — |
| KM038 | NSTE | NPSC | — | NVSY | — | NTTL | NVSK | NCSK | — | NGTY | — | NGSC |
| PE036 | NSTE | NPSC | — | NVSY | — | NTTL | NVSK | NCSK | — | NGTY | — | NGSC |
| DH084 | NSTE | NPSC | — | NVSY | — | NTTL | NVSK | NCSK | NGTY | — | NGSC | — |
| DH038 | NSTE | NPSC | — | NVSY | — | NTTL | NVSK | NCSK | NGTY | — | NGSC | — |
| NJ019 | NSTD | NPSC | — | NVSY | — | NTTL | NVSK | NCSK | NGTY | — | NGSC | — |
| NJ014 | NSTE | NPSC | — | NVSY | — | NTTL | NVSK | NCSK | NGTY | — | NGSC | — |
| BS013 | NSTE | NPSC | — | NVSY | — | NTTL | NVSK | NCSK | — | NGTY | — | NGSC |
| YX023 | NSTE | NPSC | — | NVSY | — | NTTL | NVSK | NCSK | NGTY | — | NGSC | — |
| CX023 | NSTE | NPSC | — | NVSY | — | NTTL | NVSK | NCSK | NGTY | — | NGSC | — |
| YX081 | NSTE | NPSC | — | NVSY | — | NTTL | NVSK | NCSK | — | NGTY | — | NGSC |
| CX021 | NSTE | NPSC | — | NVSY | — | NTTL | NVSK | NCSK | NGTY | — | NGSC | — |

**Table 5.**
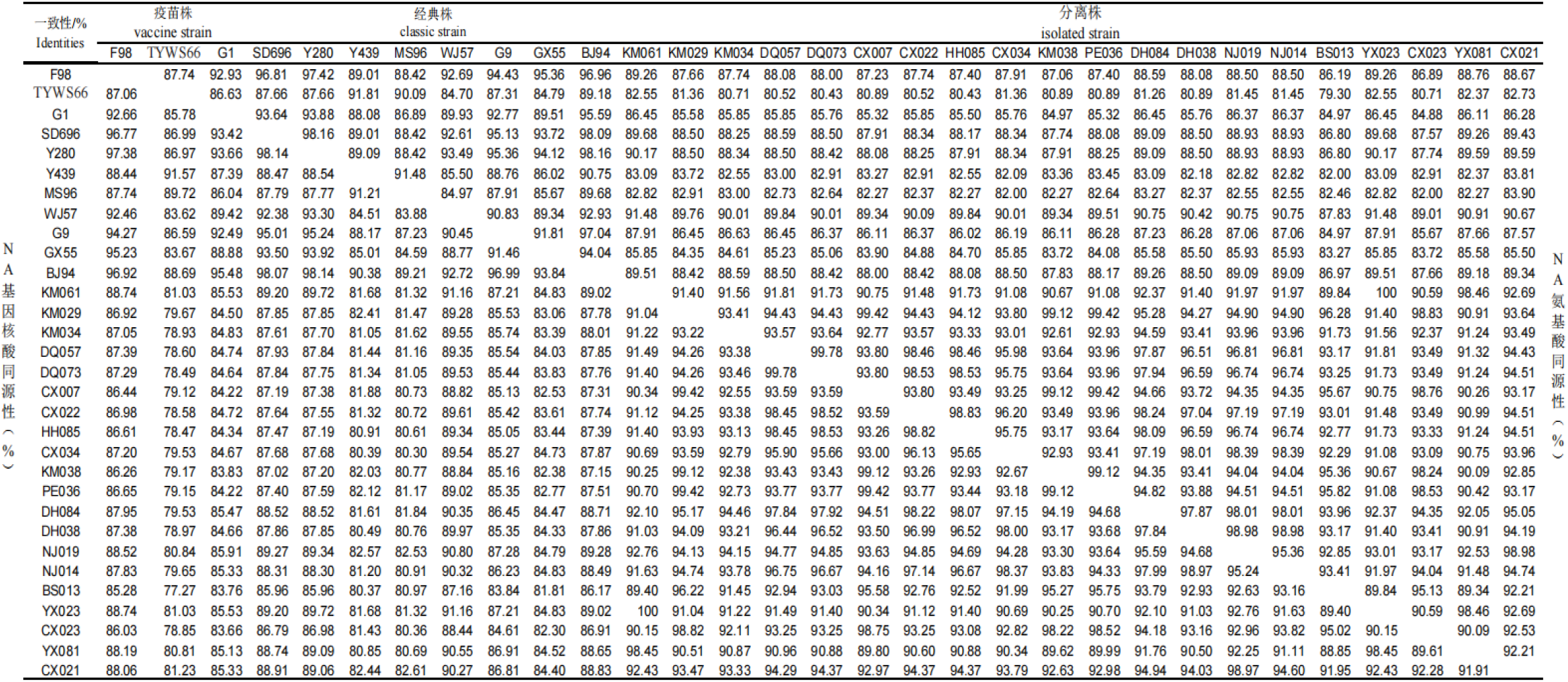
Comparison of nucleotide and amino acid sequences of NA genes of 20 H9N2 subtype AIV isolates.

### 2.2 NA gene of H9N2 avian influenza virus

#### 2.2.1 Genetic evolution and homology analysis of NA gene

The genetic evolution analysis of NA gene of 20 isolates and 11 reference strains showed that the genetic evolution tree of NA gene was divided into Eurasian lineage and North American lineage. The results showed that the NA genes of the 20 isolates belonged to the Y280 branch of the Eurasian lineage and were located on the same branch.The nucleotide and amino acid homology of NA gene of 20 isolates in Yunnan Province from 2020 to 2023 were 88.85 % - 100 % and 90.09 % -100 %, respectively. The homology within the group of isolates in 2020 was 90.50 % -98.97 % and 90.91-98.98 %, respectively. The homology within the group of isolates in 2021 was 90.69 % -100 % and 91.08 % -100 %, respectively. The homology within the group of isolates in 2022 was 92.55 % -99.78 % and 92.77 % -99.78 %, respectively. The homology within the group of isolates in 2023 was 95.02 % -99.12 % and 95.13 % -99.12 %, respectively. In addition, the nucleotide and amino acid homology of 20 isolates with vaccine strains WJ57, F98 and SD696 were 87.16 % -91.16 % and 87.83 % -91.48 %, 85.28 % -88.74 % and 86.89 % -89.26 %, 85.96 % -89.27 % and 86.8 % -91.48 %, respectively;The homology of nucleotides and amino acids between the 20 isolates and the classical strain Y280 was 85.96 % -89.72 % and 87.748 % -90.17 %, respectively, and the lowest homology with the classical TYWS66 was 77.27 % -81.23 % and 79.3 % -82.73, respectively.

#### 2.2.2 Analysis of key sites of NA gene

The complete ORF of the NA gene sequence of the 20 H9N2 subtype AIVs was composed of 1401 nucleotides and encoded 467 amino acids. All the 20 isolates lacked 9 nucleotides in the neck, encoding threonine (T), glutamic acid (E), and isoleucine (I) at positions 63,64, and 65, respectively, which belonged to the highly pathogenic NA marker and was consistent with the domestic epidemic strain represented by A/Duck/HongKong/Y280/97 strain. Hemadsorbing site (HB) was located at 366 ∼ 373,399 ∼ 404 and 431 ∼ 433, respectively. K / S/E368N, D369S/G and S/T370L mutations were detected at 366-373 sites. D399 G, N/I/S402 D/E, W/R/S403 L mutations were found in 399 ∼ 404. Q/K432R/H mutation was found at 431-433. There were no mutations in E, D, R, R, E, R, and R at 119,151,152,224,276,292, and 371 sites associated with NA protein activity and resistance to anti-influenza drugs oseltamivir and zanamivir (Table 6).

**Table 6.**
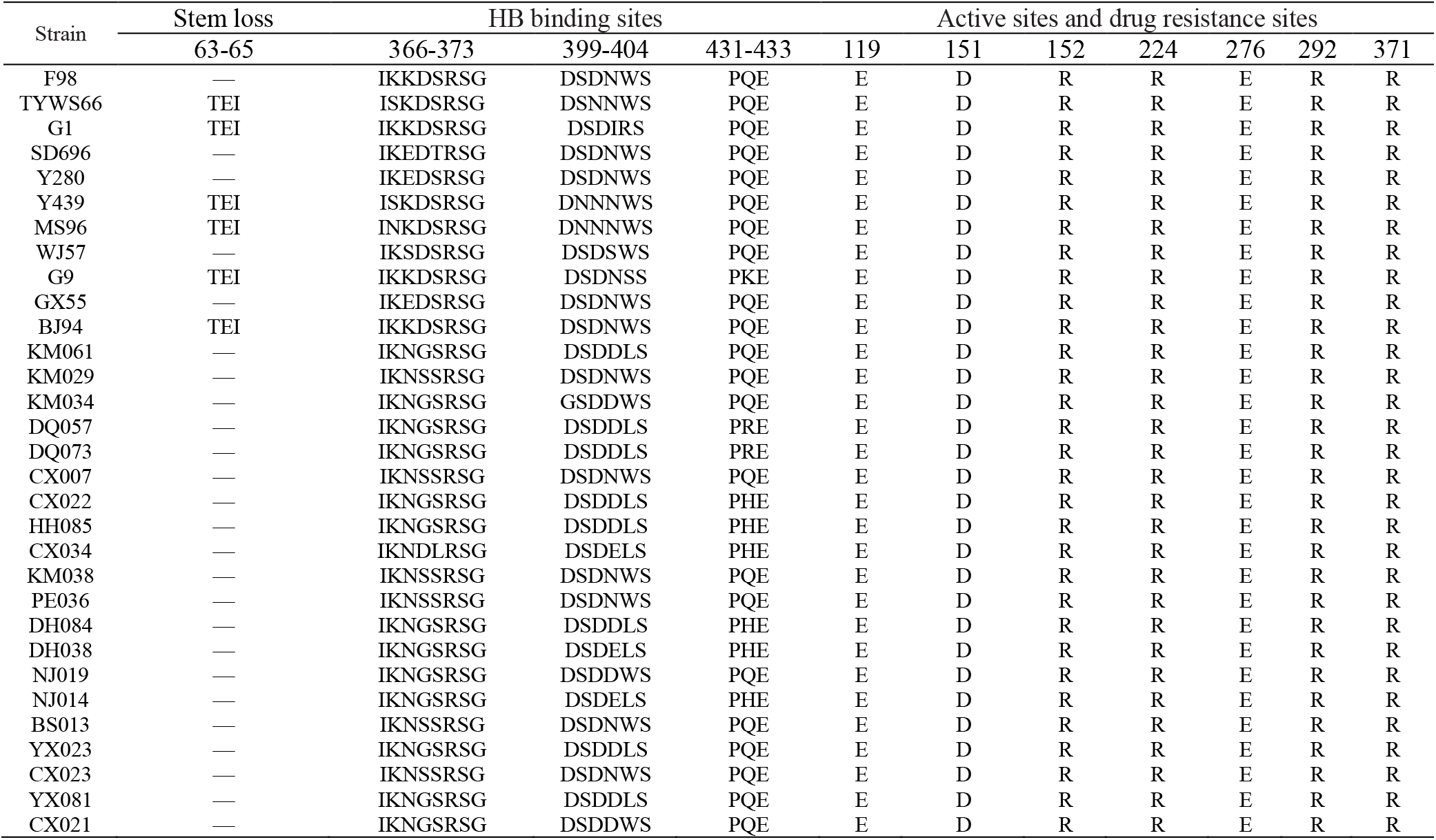
Analysis of HB binding sites, protein active sites and drug resistance sites of NA protein of 20 H9N2 subtype AIV isolates.

**Table 7.** Analysis of potential glycosylation sites of NA protein of 20 H9N2 subtype AIV isolates.

| Strain | Potential glycosylation sites of NA protein |  |  |  |  |  |  |  |  |  |  |  |  |  |  |  |  |
| --- | --- | --- | --- | --- | --- | --- | --- | --- | --- | --- | --- | --- | --- | --- | --- | --- | --- |
|  | 2 | 44 | 61 | 66 | 67 | 69 | 70 | 83 | 86 | 143 | 146 | 197 | 200 | 231 | 234 | 365 | 399 |
| F98 | — | NPSN | — | NSTT | — | — | — | NWSK | — | NGTT | — | NATA | — | NGTC | — | — | NWSG |
| TYWS66 | — | — | NITE | — | — | — | NITI | — | NWSK | — | NGTI | — | NATA | — | NGTC | — | — |
| G1 | — | — | — | — | NNTT | — | — | — | — | — | — | — | — | — | — | — | — |
| SD696 | — | NPSN | — | NSTI | — | — | — | NWSK | — | NGTT | — | NATA | — | NGTC | — | — | NWSG |
| Y280 | — | NPSN | — | NSTT | — | — | — | MWSK | — | NGTT | — | NATA | — | NGTC | — | — | NWSG |
| Y439 | — | — | NITE | — | — | NNTT | NTTI | — | NWSK | — | NGTI | — | NATA | — | NGTC | — | — |
| MS96 | — | — | NITE | — | — | NNTT | NTTI | — | NWSK | — | NGTI | — | NATA | — | NGTC | — | — |
| WJ57 | — | NPSN | — | NSTT | — | — | — | NWSK | — | NGTT | — | NATA | — | NGTC | — | — | — |
| G9 | — | NSSN | NITE | — | — | NSTT | — | — | NWSK | — | NGTA | — | NATA | — | NGTC | — | — |
| GX55 | — | — | — | NSTT | — | — | — | NWSK | — | NGTT | — | NATA | — | NGTC | — | — | NWSG |
| BJ94 | — | NPSN | NITE | — | — | NSTT | — | — | NWSK | — | NGTT | — | NATA | — | NGTC | — | — |
| KM061 | NPTQ | NSSN | — | NSTT | — | — | — | NWSK | — | NGTT | — | NATA | — | NGTC | — | NGSR | — |
| KM029 | — | NPSN | — | NSTI | — | — | — | NWSK | — | NGTT | — | NATA | — | NGTC | — | NSSR | NWSG |
| KM034 | — | NPSN | — | NNTT | NTTI | — | — | — | — | NGTT | — | NATA | — | NGTC | — | NGSR | — |
| DQ057 | — | NPSN | — | NSTT | — | — | — | NWSK | — | NGTT | — | NATA | — | NGTC | — | NGSR | — |
| DQ073 | — | NPSN | — | NSTT | — | — | — | NWSK | — | NGTT | — | NATA | — | NGTC | — | NGSR | — |
| CX007 | — | NPSN | — | NSTI | — | — | — | NWSK | — | NGTT | — | NATA | — | NGTC | — | NSSR | NWSG |
| CX022 | — | NPSN | — | NSTT | — | — | — | NWSK | — | NGTT | — | NATA | — | NGTC | — | NGSR | — |
| HH085 | — | NPSN | — | NSTT | — | — | — | NWSK | — | NGTT | — | NATA | — | NGTC | — | NGSR | — |
| CX034 | — | NPSN | — | NSTT | — | — | — | NWSK | — | NGTT | — | NATA | — | NGTC | — | — | — |
| KM038 | — | NSSN | — | NSTI | — | — | — | NWSK | — | NGTT | — | NATA | — | NGTC | — | NSRR | NWSG |
| PE036 | — | NSSN | — | NSTI | — | — | — | NWSK | — | NGTT | — | NATA | — | NGTC | — | NSSR | NWSG |
| DH084 | — | NPSN | — | NSTT | — | — | — | NWSK | — | NGTT | — | NATA | — | NGTC | — | NGSR | — |
| DH038 | — | NPSN | — | NSTT | — | — | — | — | — | NGTT | — | NATA | — | NGTC | — | NGSR | — |
| NJ019 | — | NSSN | — | NSTT | — | — | — | — | — | NGTT | — | NATA | — | NGTC | — | NGSR | — |
| NJ014 | — | NPSN | — | NSTT | — | — | — | NWSK | — | NGTT | — | NATA | — | NGTC | — | NGSR | — |
| BS013 | — | NPSN | — | NSTI | — | — | — | NWSK | — | NGTT | — | NATA | — | NGTC | — | NSSR | NWSG |
| YX023 | NPTQ | NSSN | — | NSTT | — | — | — | NWSK | — | NGTT | — | NATA | — | NGTC | — | NGSR | — |
| CX023 | — | NPSK | — | NSTI | — | — | — | NWSK | — | NGTT | — | NATA | — | NGTC | — | NSRR | NWSG |
| YX081 | — | NSSD | — | NSTT | — | — | — | NWSK | — | NGTT | — | NATA | — | NGTC | — | NGSR | — |
| CX021 | — | NSSN | — | NSTT | — | — | — | NWSK | — | NGTT | — | NATA | — | NGTC | — | NGSR | — |

The analysis of glycosylation sites showed that there were 6-8 potential glycosylation sites (Table 4) in the NA protein of 20 isolates. Compared with the reference strain, except for 2021, there was an additional glycosylation site (NWSG) at position 399 in other years. In addition, the glycosylation site 44 of the isolates YX018 and CX020 in 2020 was mutated from NPSN to NSSD and NSSN, and the NSSR at glycosylation site 365 of the isolate KM029 was inconsistent with other isolates. In 2021, the glycosylation site 2 of the isolates KM061 and YX023 increased by one NPTQ, and the glycosylation sites 83 and 365 of the isolates NJ019 and CX034 were missing; in 2022, the glycosylation site 66 of the isolate KM034 had a mutation from NSTT (I) to NNTT, a glycosylation site NTTI was added at position 67, and a glycosylation site was missing at position 83. The isolate CX007 had one additional site at glycosylation sites 365 and 399, respectively, which were NSSR and NWSG. The glycosylation site 365 of KM038 and PE036 isolates in 2023 increased by one glycosylation site, NSRR and NSSR, respectively. The NP (S) SN to NPSK mutation occurred at position 44 of the glycosylation site of the isolate CX023, and one glycosylation site was added at position 365, which was NSRR.

## 3 Discussions

H9N2 subtype avian influenza virus (AIV) is widely spread throughout China, which not only has a negative impact on poultry production performance, but also has the potential risk of cross-species transmission to infect humans, posing a serious threat to the healthy development of China ‘s poultry industry and public health security. The H9N2 avian influenza virus is constantly mutating under the influence of environmental and natural selection pressures. As the virus evolves, the host range is gradually expanding. It can not only infect poultry, but also infect pigs and mammals such as humans [7], indicating that the H9N2 subtype avian influenza virus has a hidden infection in animals and has the possibility of causing public health problems.

The HA and NA genes of H9N2 subtype AIV are divided into two branches : Eurasian and North American. The Eurasian branch can be divided into three branches: Y280(BJ/94) - like, G1-like and Y439-like[8]. After the A / Duck / Hong Kong / Y280 / 97 strain was reported in 1997, the strain represented by Y280-like became the dominant type in China[9]. The 20 isolates isolated in Yunnan Province in this study were closely related to the Y280-like branch in the Eurasian branch, belonging to the same branch, which was consistent with the national epidemic situation. The homology of HA genes among the 20 isolates in this study was relatively close. The nucleotide and amino acid homology were 88.46 % ∼ 99.81 % and 89.08 % ~ 99.61 %, respectively. The nucleotide and amino acid homology of NA gene were 88.85 % ~ 100 % and 90.09 % ∼ 100 %, respectively.This shows that HA and NA genes change over time, and mutations are constantly accumulating. In this study, the HA gene similarity of 20 isolates was lower than that of 3 vaccine strains, indicating that the H9N2 avian influenza virus epidemic isolates in Yunnan Province were constantly mutating under the pressure of host immunity. At the same time, the homology between virus strains in different regions was higher, but the earlier viruses had different degrees of variation and evolution.

The changes of important amino acid sites in HA protein affect the biological function of avian influenza virus[10]. The amino acid sequence of HA protein cleavage site is the key factor to determine the pathogenicity and virulence of avian influenza virus[11]. The results of this study showed that the HA protein cleavage site sequence of 20 isolates was PSRSSRGLF, and there were continuous alkaline amino acids, showing the characteristics of low pathogenic avian influenza, which was consistent with the characteristics of H9 low pathogenic avian influenza strains prevalent in recent years. The amino acid composition of the receptor binding site of the HA protein affects the receptor binding characteristics and host range of the virus[12]. Studies have shown that when 226 (H9 sequence at position 234) is glutamic acid (Q), the avian influenza virus mainly binds to the receptor α-2,3-galactose-linked sialic acid. When 226 is leucine D (L), the human influenza virus mainly binds to the receptor α-2,6-galactose-linked sialic acid[13]. In the absence of leucine (L) at position 234 (H9 number). The 163T or 191N substitution may lead to the binding ability of some H9 viruses to obtain α2-6-SA [14, 15]. In this study, the 234th site of the 20 isolates was L, and the 191st site of some isolates was N and 163 was T. The changes of these sites make them have the ability to bind to mammalian sialic acid α2-6 receptors and have the tropism of human infection. The Q234 L mutation indicates that the H9N2 subtype avian influenza virus circulating in the live poultry market in Yunnan Province may infect humans. Therefore, it is necessary to strengthen the monitoring of H9N2 subtype avian influenza virus variation in Yunnan Province. The antigenicity and virulence of HA protein glycosylation sites are affected by the number and location of glycosylation sites. The increase of glycosylation sites near the HA cleavage site will prevent protease cleavage and virus infection, while the glycosylation sites near the receptor binding site will affect the binding level of the virus to the host cell [16]. In this study, 20 isolates all lacked one glycosylation site at position 218 of HA gene. Four isolates in 2020, five in 2021, two in 2022 and one in 2023 increased one glycosylation site at positions 313, 491 and 550, respectively. The appearance of glycosylation sites may change their biological characteristics and infect new hosts. In addition, the increased 313 glycosylation site is close to the HA cleavage site, which will enhance the binding ability of the virus to the receptor, and enhance the ability of the virus to infect cells and replicate in cells[17, 18].

In this study, we compared the antigenicity-related sites of reference strains and isolates, and found that there were 6 mutations in the antigenic sites of 7 HA of 20 isolates from 2020 to 2023, indicating that the antigenicity of H9 subtype AIV prevalent in Yunnan Province has undergone deep variation. It also suggests that immune pressure can promote the mutation of strains. Studies have shown that the mutation of the 168 th amino acid from aspartic acid (D) to aspartic acid (N) may lead to antigen drift of H9 subtype avian influenza virus to evade vaccine immunization[19]. In this study, the 168 antigen site of 13 isolates was N, indicating that the H9 subtype AIV in Yunnan Province had vaccine immune evasion from 2020 to 2023, which may lead to the failure of the current H9N2 subtype AIV vaccine immunization. In addition, there are 6 strains with 168 antigen sites E, and its biological characteristics need further study.

NA protein plays an important role in viral replication and escape from cells. The NA protein head has an enzyme activity center and an antigenic site[20]. The head enzyme activity center, transmembrane domain and cytoplasmic domain of NA protein are connected by the neck. The length of the neck is related to the cross-species transmission ability and pathogenicity of the virus[21]. The deletion of the neck of the NA protein is a sign of increased virulence of the H9N2 subtype avian influenza virus[22]. In this study, the deletion of TEI 3 amino acids occurred in the 62-64 amino acid region of NA protein stem of all isolates from 2020 to 2023, which enhanced the activity of NA protease and the ability to release from cells, which was consistent with the domestic epidemic strain represented by A / Duck / Hong Kong / Y280 / 97 strain. Under normal circumstances, the HB site of NA gene is 366-373,399-404 and 431-433 amino acids forming three circular structures. Amino acids and sialic acid are directly bound to the formed circular structure. The amino acid mutation of HB will affect the replication and growth of the virus[23]. K/S/E368N, D369S/G and S/T370L mutations were found in 366 ∼ 373 sites of some isolates in this study. D399 G, N/I/S402D/E, W/R/S403 L mutations were found in 399 ∼ 404. Q / K432R / H mutations are produced at 431-433 sites. These mutations may regulate the binding of sodium enzyme activity to HA receptor to achieve the best results, so that the virus can proliferate and grow better. Studies have reported that influenza virus NA protease activity site or key amino acid site specific mutations: E119A/ D/G/V, D151E, R152K, R224K, E276D, R292K, R371K, the virus will enhance the resistance to neuraminidase inhibitor drugs (oseltamivir, zanamivir)[24, 25]. In this study, there were no mutations in the NA protease active site and key site amino acids of the 20 strains, indicating that these isolates were not resistant to neuraminidase inhibitors. Glycosylation sites are important for maintaining the structure and normal biological function of NA protein molecules. Glycosylation modification can regulate neuraminidase activity and is related to the neurotropism of influenza virus[26]. There were 6-8 potential glycosylation sites in the NA gene of the isolates in this study. Compared with other isolates, KM061 and YX023 had a potential glycosylation site NPTQ at the second position, and KM034 added a potential glycosylation site NTTI at the 67 th position. Its biological characteristics need further study. Studies have shown that the glycosylation at position 44 can affect the activity and virulence of NA, and the increase of the glycosylation site at position 44 will enhance the replication rate of the virus in MDCK and A549 cells[27]. In this study, all 20 isolates had a glycosylation site at position 44, but compared with the classical strain and vaccine strain, the 44th glycosylation site of the isolates CX023 and YX081 had NPSK and NSSD mutations. Whether the mutation has an effect on the proliferation of the virus needs further study. The deletion of the 146th glycosylation site of the NA protein changes the steric hindrance of the N protein and the presence of Lys residues at the C-terminus enhances viral pathogenicity[28]. In this study, there was a deletion of 146 glycosylation site in all 20 isolates, but there was no Lys residue at the C-terminus, and its virulence needs to be further studied. The glycosylation site at position 402 is the sialic acid binding site of NA gene, which is located in the antigenic determinant region. The increase or deletion of this glycosylation site may change the enzyme activity and antigenicity of H9N2 subtype avian influenza and affect its pathogenicity.

## 4. Conclusion

The results of this study show that the HA and NA genes of H9N2 subtype avian influenza virus in Yunnan Province have evolved continuously, but they still belong to the Y280 branch of the Eurasian lineage. The cleavage site of HA protein of 20 isolates was 333-341, the sequence was PSRSSRGLF, containing continuous basic amino acids, which was consistent with the molecular characteristics of low pathogenic avian influenza virus. The HA receptor binding sites were L, N and T at positions 234,191 and 163. The changes of these sites make it have the ability to bind to mammalian sialic acid α-2,6 sialic acid receptors and have the tropism to infect humans. There are multi-point variations in the antigenic sites of HA, and the variation of the antigenic sites reduces the matching with the vaccine strain, thereby reducing the protective effect of the vaccine and increasing the risk of infection. NA protein neck deletion of 62,63,64 amino acids (TEI), which is a highly pathogenic avian influenza marker; There are 6-8 potential glycosylation sites in the NA protein, and there are 3 new glycosylation sites. The emergence of new glycosylation sites may change the antigenicity of the virus and affect the receptor binding characteristics to infect new hosts. There were no mutations in the amino acids at the active sites and key sites of NA protease, suggesting that the isolates in Yunnan Province did not produce resistance to neuraminidase inhibitors. There are many variations in the 366 ∼ 373,399 ∼ 404 and 431 ∼ 433 amino acids of the red blood cell binding (HB) site of NA protein, which enables the virus to proliferate and grow better in host cells.

In summary, the receptor sites and glycosylation sites of H9N2 subtype avian influenza virus in the live poultry market of Yunnan Province are diversified, and the mutations of some key sites may cause the enhancement of infectivity and transmission. The monitoring of low pathogenic H9N2 subtype avian influenza virus should be strengthened to prevent it from breaking the interspecies barrier and spreading to humans and lower mammals, so as to prevent the occurrence of H9N2 subtype avian influenza epidemic.

## References

1. Guo YJ, Krauss S, Senne DA, Mo IP, Lo KS, Xiong XP, Norwood M, Shortridge KF, Webster RG, Guan Y: Characterization of the pathogenicity of members of the newly established H9N2 influenza virus lineages in Asia. Virology 2000, 267:279–288.

2. Shen HQ, Yan ZQ, Zeng FG, Liao CT, Zhou QF, Qin JP, Xie QM, Bi YZ, Chen F: Isolation and phylogenetic analysis of hemagglutinin gene of H9N2 influenza viruses from chickens in South China from 2012 to 2013. J Vet Sci 2015, 16:317–324.

3. Yuanji G JLIX: Discovery of men infected by avian influenza A(H9N2) virus. Chinese Journal of Experimental and Clinical Virology 1999, 13:1052–1081.

4. Lam TT, Wang J, Shen Y, Zhou B, Duan L, Cheung CL, Ma C, Lycett SJ, Leung CY, Chen X, et al: The genesis and source of the H7N9 influenza viruses causing human infections in China. Nature 2013, 502:241–244.

5. Fouchier RA, Munster V, Wallensten A, Bestebroer TM, Herfst S, Smith D, Rimmelzwaan GF, Olsen B, Osterhaus AD: Characterization of a novel influenza A virus hemagglutinin subtype (H16) obtained from black-headed gulls. J Virol 2005, 79:2814–2822.

6. Xing Z, Cardona CJ, Li J, Dao N, Tran T, Andrada J: Modulation of the immune responses in chickens by low-pathogenicity avian influenza virus H9N2. J Gen Virol 2008, 89:1288–1299.

7. Choi YK, Ozaki H, Webby RJ, Webster RG, Peiris JS, Poon L, Butt C, Leung YH, Guan Y: Continuing evolution of H9N2 influenza viruses in Southeastern China. J Virol 2004, 78:8609–8614.

8. Zhang Y, Yin Y, Bi Y, Wang S, Xu S, Wang J, Zhou S, Sun T, Yoon KJ: Molecular and antigenic characterization of H9N2 avian influenza virus isolates from chicken flocks between 1998 and 2007 in China. Vet Microbiol 2012, 156:285–293.

9. Liu Q, Zhao L, Guo Y, Zhao Y, Li Y, Chen N, Lu Y, Yu M, Deng L, Ping J: Antigenic Evolution Characteristics and Immunological Evaluation of H9N2 Avian Influenza Viruses from 1994-2019 in China. Viruses 2022, 14.

10. Jin X, Zha Y, Hu J, Li X, Chen J, Xie S, Dai Y, Li Z, Wang X, Wang F, et al: New molecular evolutionary characteristics of H9N2 avian influenza virus in Guangdong Province, China. Infect Genet Evol 2020, 77:104064.

11. Hutter J, Rodig JV, Hoper D, Seeberger PH, Reichl U, Rapp E, Lepenies B: Toward animal cell culture-based influenza vaccine design: viral hemagglutinin N-glycosylation markedly impacts immunogenicity. J Immunol 2013, 190:220–230.

12. Dortmans JC, Dekkers J, Wickramasinghe IN, Verheije MH, Rottier PJ, van Kuppeveld FJ, de Vries E, de Haan CA: Adaptation of novel H7N9 influenza A virus to human receptors. Sci Rep 2013, 3:3058.

13. Wan H, Perez DR: Amino acid 226 in the hemagglutinin of H9N2 influenza viruses determines cell tropism and replication in human airway epithelial cells. J Virol 2007, 81:5181–5191.

14. Bi Y, Li J, Li S, Fu G, Jin T, Zhang C, Yang Y, Ma Z, Tian W, Li J, et al: Dominant subtype switch in avian influenza viruses during 2016-2019 in China. Nat Commun 2020, 11:5909.

15. Li X, Shi J, Guo J, Deng G, Zhang Q, Wang J, He X, Wang K, Chen J, Li Y, et al: Genetics, receptor binding property, and transmissibility in mammals of naturally isolated H9N2 Avian Influenza viruses. Plos Pathog 2014, 10:e1004508.

16. Liu M, Chen H, Luo F, Li P, Pan Q, Xia B, Qi Z, Ho WZ, Zhang XL: Deletion of N-glycosylation sites of hepatitis C virus envelope protein E1 enhances specific cellular and humoral immune responses. Vaccine 2007, 25:6572–6580.

17. Tsuchiya E, Sugawara K, Hongo S, Matsuzaki Y, Muraki Y, Nakamura K: Role of overlapping glycosylation sequons in antigenic properties, intracellular transport and biological activities of influenza A/H2N2 virus haemagglutinin. J Gen Virol 2002, 83:3067–3074.

18. Peng Q, Zhu R, Wang X, Shi H, Bellefleur M, Wang S, Liu X: Impact of the variations in potential glycosylation sites of the hemagglutinin of H9N2 influenza virus. Virus Genes 2019, 55:182–190.

19. Okamatsu M, Sakoda Y, Kishida N, Isoda N, Kida H: Antigenic structure of the hemagglutinin of H9N2 influenza viruses. Arch Virol 2008, 153:2189–2195.

20. Lamb RA, Choppin PW: The gene structure and replication of influenza virus. Annu Rev Biochem 1983, 52:467–506.

21. Yang F, Xiao Y, Liu F, Yao H, Wu N, Wu H: Molecular characterization and antigenic analysis of reassortant H9N2 subtype avian influenza viruses in Eastern China in 2016. Virus Res 2021, 306:198577.

22. Spackman E: A Brief Introduction to Avian Influenza Virus. Methods Mol Biol 2020, 2123:83–92.

23. Smith GL, Hay AJ: Replication of the influenza virus genome. Virology 1982, 118:96–108.

24. Yen HL HETG: Importance of neuraminidase active-site residues to the neuraminidase inhibitor resistance of influenza viruses. J Virol 2006, 80:8787–8795.

25. Hurt AC, Holien JK, Parker M, Kelso A, Barr IG: Zanamivir-resistant influenza viruses with a novel neuraminidase mutation. J Virol 2009, 83:10366–10373.

26. Skehel JJ, Stevens DJ, Daniels RS, Douglas AR, Knossow M, Wilson IA, Wiley DC: A carbohydrate side chain on hemagglutinins of Hong Kong influenza viruses inhibits recognition by a monoclonal antibody. Proc Natl Acad Sci U S A 1984, 81:1779–1783.

27. Wu CY, Lin CW, Tsai TI, Lee CD, Chuang HY, Chen JB, Tsai MH, Chen BR, Lo PW, Liu CP, et al: Influenza A surface glycosylation and vaccine design. Proc Natl Acad Sci U S A 2017, 114:280–285.

28. Goto H, Kawaoka Y: A novel mechanism for the acquisition of virulence by a human influenza A virus. Proc Natl Acad Sci U S A 1998, 95:10224–10228.

